# Polychrome activity profiling distinguishes Cys proteases and their isoforms in plants

**DOI:** 10.64898/2026.06.15.732375

**Authors:** Kaijie Zheng, Mariana Schuster, Nattapong Sanguankiattichai, Till Kessenbrock, Markus Kaiser, Renier A. L. van der Hoorn

**Affiliations:** Key Laboratory of Soybean Molecular Design Breeding, Northeast Institute of Geography and Agroecology, Chinese Academy of Sciences, Changchun 130102, China; The Plant Chemetics Laboratory, Department of Biology, University of Oxford, OX1 3RB Oxford; Department of Microbiology, Faculty of Science, Mahidol University; Bangkok, Thailand; ZMB Chemical Biology, Faculty of Biology, University of Duisburg-Essen, Essen, Germany

**Keywords:** protease isoforms, papain-like protease, fluorescent labeling, activity-based protein profiling

## Abstract

Activity-based profiling with fluorescent probes is a powerful tool for the functional characterization of whole enzyme classes in crude proteomes. Here, we discovered that probe cocktails consisting of the E-64 warhead carrying different fluorophores via short linkers distinguishes the labeling of papain-like cysteine proteases and their isoforms because of their differential affinity to these probes. This causes ‘polychrome labeling’ with specific apparent colors for different protease families and isoforms. Polychrome labeling revealed differential labeling of isoforms of RD21-like proteases carrying C-terminal granulin domains. Molecular modeling of the RD21 isoforms revealed that the C-terminal granulin may create a fluorophore-binding pocket with the substrate-binding groove, implying that the granulin domain influences substrate selectivity. The concept of polychrome labeling may be widely applicable to other chemical probes for the characterization of protein families in all kingdoms of life.

## INTRODUCTION

Activity-based probes are important tools in the functional characterization of proteins in proteomes. Activity-based probes for proteases, lipases, glycosidases, cytochrome P450s, kinases, phosphatases and many other proteins have uncovered protein activities under specific conditions and have been used to discover and optimize inhibitors of these proteins (Morimoto & Van der Hoorn, 2016; Chakrabarty et al., 2019; Niphakis et al., 2024).

Detection and identification of labeled proteins is mostly achieved by fluorescence or by mass spectrometry, respectively (Kovacs & Van der Hoorn, 2016). Different fluorophores are used for the detection of labeled proteins separated on protein gels. Mixing of proteomes labeled with different fluorophores has facilitated the direct comparison of labeling between proteomes on 1D and 2D gels. Difference gel electrophoresis (DIGE), for instance, was used to detect selective inhibition of cathepsins in liver proteomes (Greenbaum et al., 2002), to compare in-extract with in-cell labeling of serine hydrolases in human cells (Gillet et al., 2008), and to detect differential serine hydrolase activities in plants upon infection (Hong & Van der Hoorn, 2016). DIGE works under the assumption that the different fluorophores do not influence labeling.

Fluorescent probes that target different proteins can also be mixed to simultaneously label different enzyme classes and distinguish them using the different fluorophores. Such approach displayed different serine proteases in neutrophils using fluorescent probes that contain selective recognition sequences (Kasperkiewicz et al., 2017), and displayed different catalytic subunits of the proteasome using a cocktail of different fluorescent probes in mammals and plants (De Bruin et al., 2016; Misas-Villamil et al., 2017). Likewise, cocktails of fluorescent probes targeting different enzymes were used to detect differentially active α-glucosidases and serine hydrolases in crocus upon infection with *Fusarium oxysporum* (Husaini et al., 2018).

One class of proteins studied with activity-based probes are papain-like cysteine proteases (PLCPs) from MEROPS family C1A in clan CA (Rawlings et al., 2018). PLCPs are produced by most kingdoms of life. Plants, for instance, produce over 25 PLCPs that belong to nine subfamilies (Richau et al., 2012). In animals, PLCPs are better known as cathepsins (Fonovic & Turk, 2014). PLCPs fold in two lobes that carry a catalytic triad with the catalytic cysteine in the middle of the substrate groove created in-between the two lobes. PLCPs are produced with an autoinhibitory prodomain and some carry C-terminal fusions with other domains, such as the granulin domain. Most PLCPs are irreversibly and covalently inhibited by E-64, a natural dipeptide-like epoxide that reacts with the catalytic Cys residue. DCG-04 is a biotinylated derivative of E-64 that has been frequently used to identify active PLCPs in proteomes by mass spectrometry (Greenbaum et al., 2000; Kocks et al., 2003; Van der Hoorn et al., 2004; Hook et al., 2012; Schulze-Hunck et al., 2019). Also fluorescent derivatives of E-64 have been used to label and detect active PLCPs in proteomes (Greenbaum et al., 2002; Shabab et al., 2008; Richau et al., 2012; Passarge et al., 2021).

Here, we generated and tested fluorescent E-64 derivatives with short linkers by coupling a commercially available E-64 derivative carrying an azide to three different Bodipy fluorophores carrying an alkyne. We found that these three probes do not label PLCPs with equal affinity and discovered that labeling with probe cocktails results in polychrome labeling, where PLCPs and their isoforms can be distinguished by different apparent colors caused by labeling with specific probe ratios.

## RESULTS AND DISCUSSION

To produce fluorescent activity-based probes for PLCPs, we performed copper-catalysed cycloaddition reactions (click chemistry) to couple commercially available azide-E-64 with three different alkyne-labeled Bodipy fluorophores, resulting in TK009 carrying BDP-630/50, TK010 carrying BDP-TMR, and TK011 carrying BDP-FL, which can be detected using Cy5, Cy3 and Cy2 settings, respectively (**Figure 1**).

**Figure 1.**
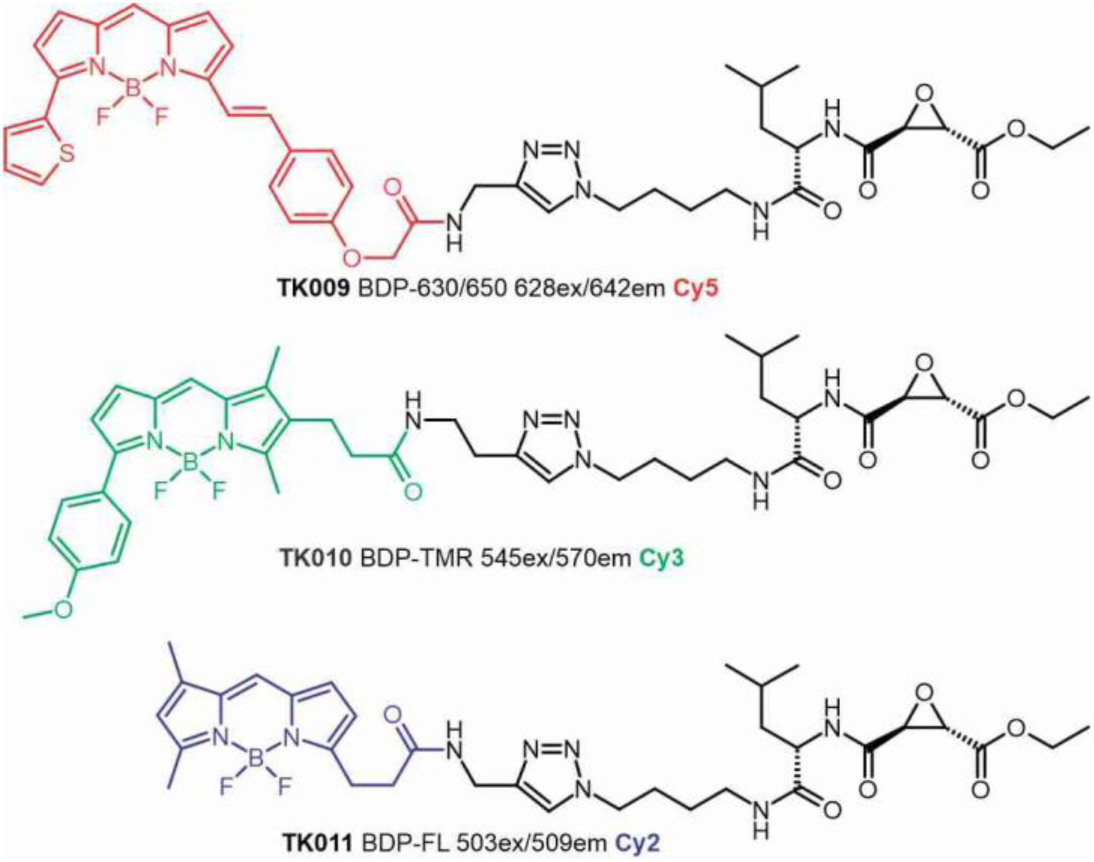
Structure of the polychrome probes. Probes carry the E-64 warhead, coupled to three different Bodipy fluorophores via a click chemistry triazole linker.

To test the probes, we transiently expressed Arabidopsis RD21, a well-described PLCP, by agroinfiltration of *Nicotiana benthamiana* (Gu et al., 2012). RD21 is known to exist in multiple active isoforms that differ in the C-terminal extension (**Figure 2A**). Intermediate RD21 (iRD21) is 35 kDa and carries the granulin domain at the C-terminus, whilst 22-25 kDa mature RD21 (mRD21) only consists of the catalytic domain with various small C-terminal extensions caused by differential processing (Gu et al., 2012). Both iRD21 and mRD21 are active proteases since they lack the N-terminal autoinhibitory prodomain which is removed by yet-unidentified endogenous plant proteases (Gu et al., 2012).

**Figure 2.**
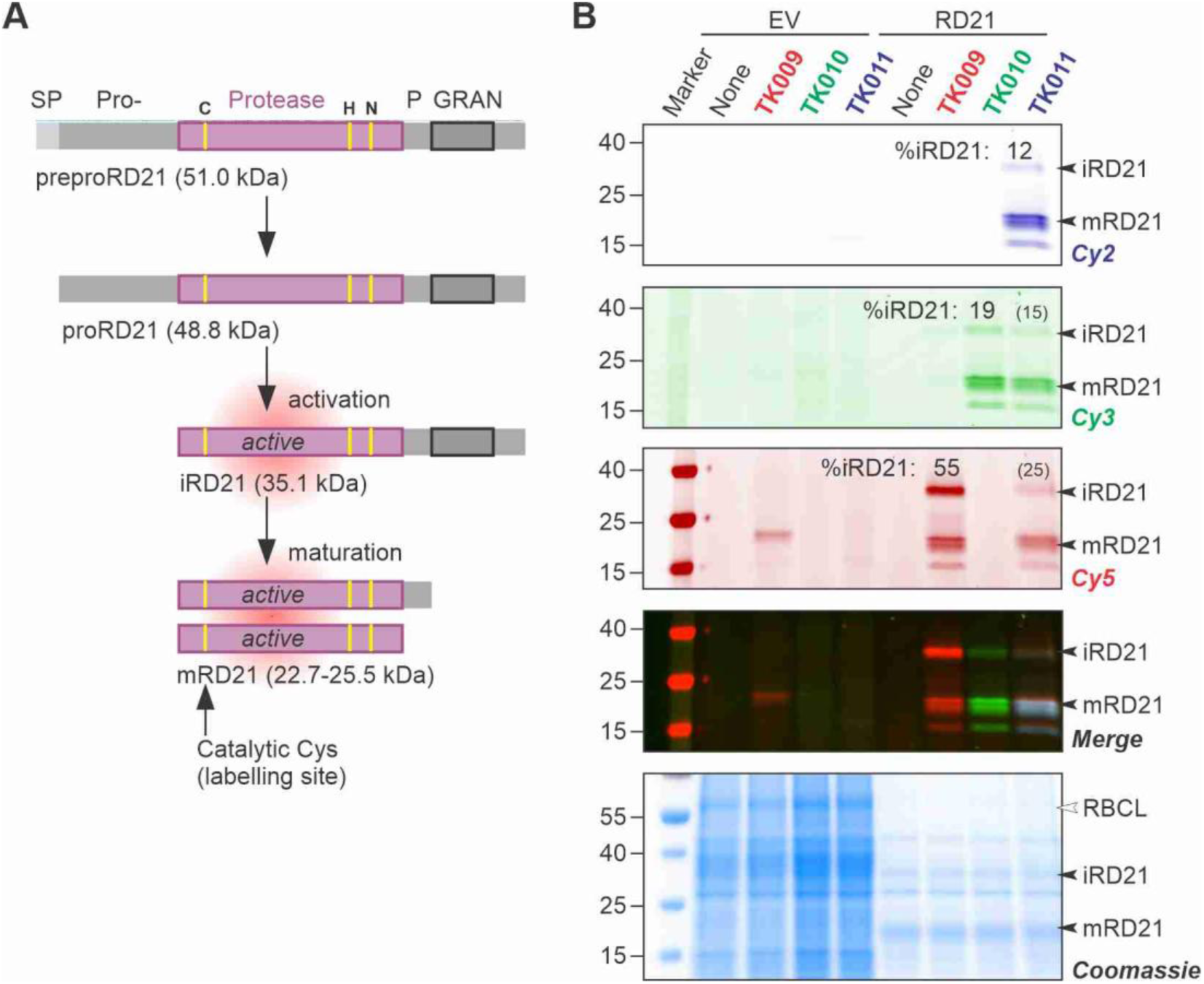
RD21 is labeled by all three probes. **(A)** Conversion of the initial translation product into active intermediate and mature RD21. **(B)** Total extracts of leaves expressing RD21 or the empty vector (EV) control) were incubated with/out 1 µM TK009, TK010 and TK011 for 3 hours, separated on gel and scanned for fluorescence and stained with Coomassie. Indicated are the intermediate (i) and mature (m) isoforms of RD21 and the large subunit of ribulose-1,5-bisphosphate carboxylase/oxygenase (RBCL). This experiment was repeated three times with a similar outcome.

TK009 labeled active RD21 isoforms causing specific signals in the Cy3 channel and not in the Cy2 and Cy5 channel and these signals were mostly absent from the empty vector (EV) control and absent from the no-probe control (**Figure 2B**). TK010 labeled the same active RD21 isoforms and was detected in the Cy5 channel but not in the Cy2 or Cy3 channels (**Figure 2B**). A weaker signal was detected with TK010 in the EV control that probably represents endogenous *N. benthamiana* PLCP(s). TK011 labeled the same active RD21 isoforms and is detected in the Cy2 channel, but in this case the fluorophore was also detected in the Cy3 and Cy5 channels but with reduced intensity when compared to TK010 and TK009, respectively (**Figure 2B**). The merged picture demonstrates that all three probes label the same RD21 isoforms, but with different efficiencies: TK009 labeled iRD21 more than mRD21 isoforms, whereas TK010 and TK011 label mRD21 more than iRD21 isoforms. In other words, 55% of the TK009 signal is from iRD21, whereas iRD21 comprises 19% and 12% of the total signal from TK010 and TK011, respectively (**Figure 2B**). Please note that the incubation of extracts of RD21 expressing leaves for 3hrs at room temperature resulted in a nearly complete degradation of the total proteome, with the exception of proteins that migrate at similar molecular weights as the labeled proteins, suggesting that RD21 degrades everything except itself (**Figure 2B**).

To confirm the differential labeling, we mixed the three labeled RD21 samples together (‘label-and-mix’) and separated this mixed sample on gel. This confirmed the different labeling efficiencies, resulting in a red iRD21 signal, and light green mRD21 signals (**Figure 3**). We also performed an RD21 labeling with a cocktail of the three probes mixed together (‘mix-and-label’). Remarkably, however, competitive labeling of RD21 with the three probes simultaneously causes more mRD21 labeling by TK011, whilst reducing mRD21 labeling by TK009 and TK010 (**Figure 3**). Thus, when mixed together, TK011 is able to label mRD21 faster than TK009 or TK010.

**Figure 3.**
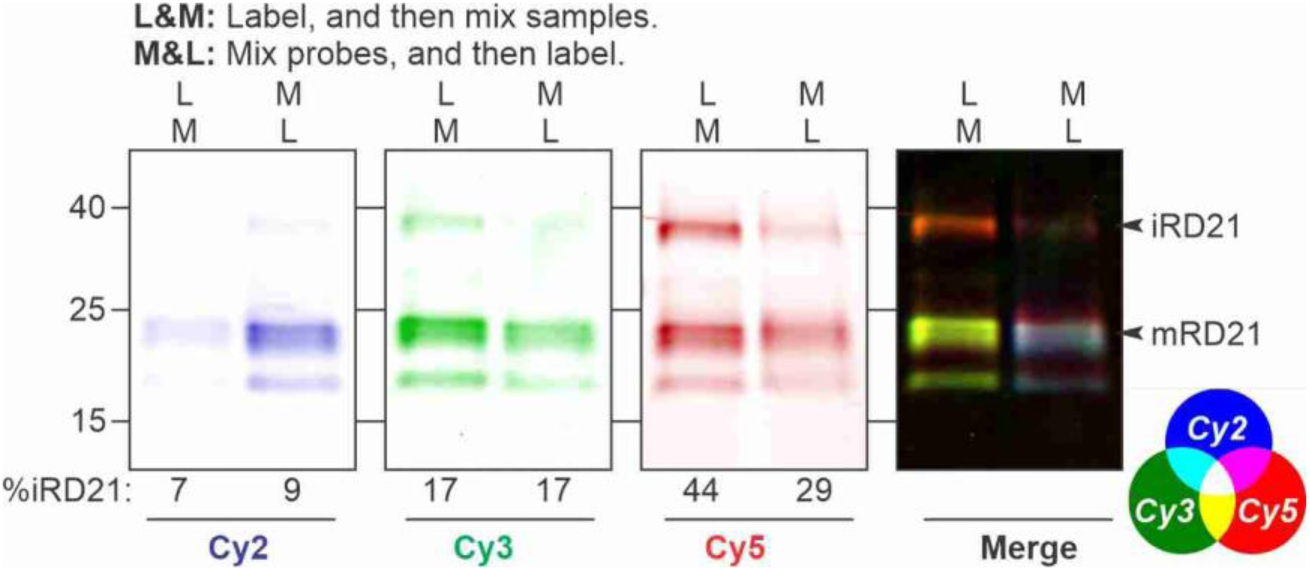
Labeling ratio is different upon competitive labeling. Extracts of leaves transiently expressing RD21 were labeled with 1 µM TK009, TK010 and TK011 for 3 hours and then mixed (L&M), or labeled with a cocktail containing 1 µM TK009, TK010 and TK011 for 3 hours (M&L). Proteins were separated on protein gel and scanned for fluorescence. This experiment was repeated three times with a similar outcome. The Coomassie-stained gel is shown in Supplemental **Figure S3**.

We next performed various experiments to test if competitive labeling of RD21 is dependent on conditions. First, in time course experiments, all three probes reach maximal labeling after one hour with indistinguishable kinetics (**Figure 4A**, Supplemental **Figure S1A**), suggesting that free probe ratios remain similar during the labeling time course. Second, labeling at various pH revealed that RD21 labeling occurs at pH 4-8 but there is an optimum at pH 6.5 and another optimum at pH 5.0 (**Figure 4B**, Supplemental **Figure 1B**), which is consistent with previous labeling experiments (Richau et al., 2012). All three probes show this pH-dependent labeling pattern but labeling at lower pH was less efficient for TK009 when compared to the other two probes (**Figure 4B**, Supplemental **Figure 1B**). Third, labeling at various probe cocktail concentrations revealed that signal saturation is reached at 0.2 µM and that TK009 labeling is relatively weak at 0.2 µM and lower probe concentrations (**Figure 4C**, Supplemental **Figure 2A**), indicating that it has less competitive strength compared to the other two probes. Finally, labeling at various protein concentrations revealed a linear correlation between signal intensity and protein concentration and this correlation is indistinguishable for the three probes (**Figure 4D**, Supplemental **Figure S2B**). Taken together, these experiments indicate that the labeling ratio between the probes does not depend on time or protein concentration as long as the probes are used in excess. Limited probe concentrations and a different pH may shift the ratio slightly. At standard labeling conditions (0.2 µg/ml protein for 3 hours at pH 6.0 with 1 µM TK probe cocktail) the probe excess causes robust polychrome labeling.

**Figure 4.**
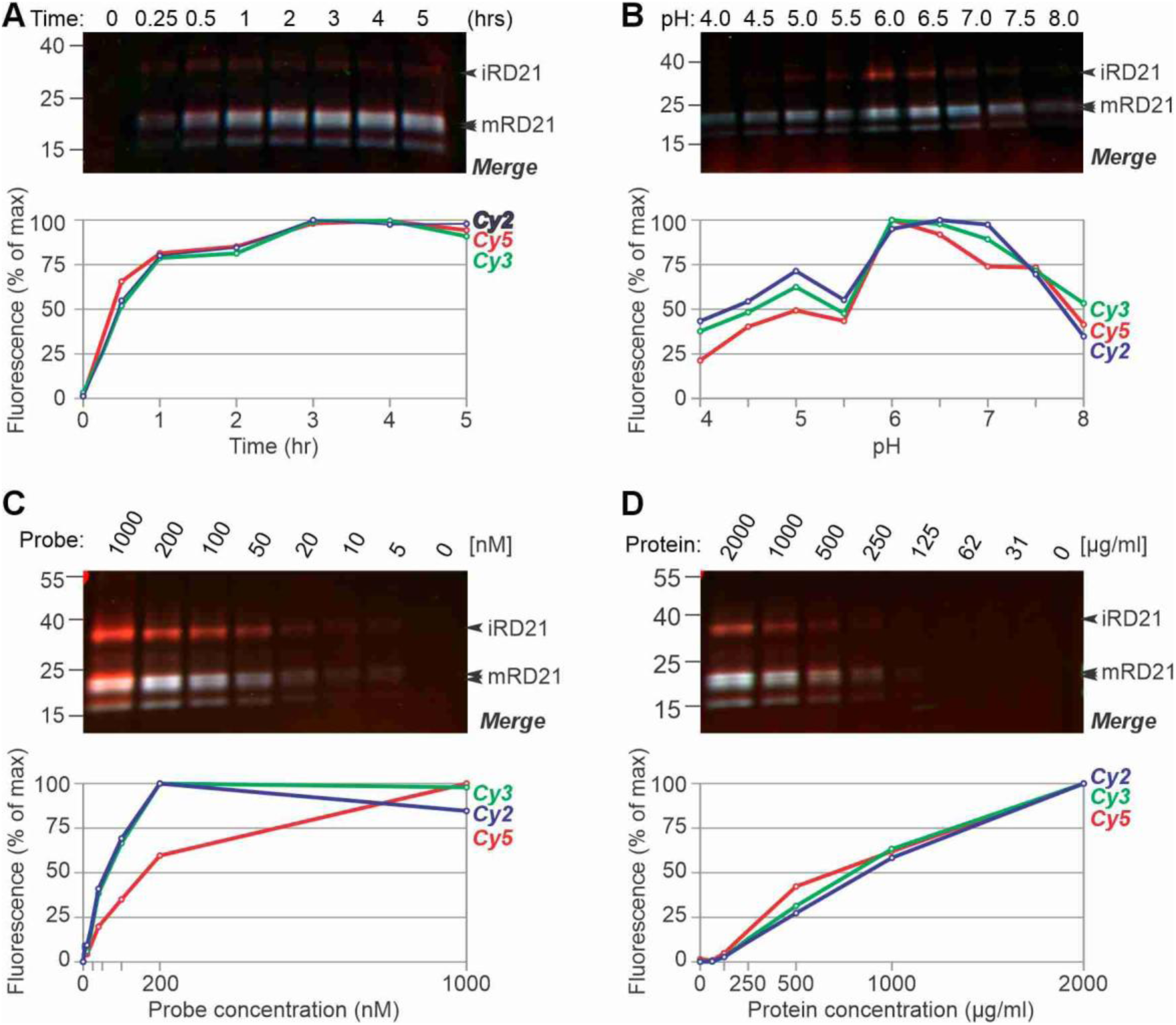
Polychrome RD21 labeling at various conditions. Total extracts of agroinfiltrated leaves expressing RD21 were incubated with for various times **(A)**, at various pH **(B)**, and at various probe **(C)** and protein **(D)** concentrations. Standard conditions were 1µM TK probe cocktail at pH 6.0 for 3 hours. Samples were precipitated in acetone and separated on protein gels, scanned for fluorescence and stained with Coomassie. All fluorescent signal intensities were quantified and plotted as a percentage of the strongest signal for each fluorescent signal. This experiment was repeated with a similar outcome. The Coomassie-stained gel and the split channels are shown in Supplemental **Figures S1 and S2**.

The differential labeling efficiencies of the probes on the different RD21 isoforms was unexpected because they carry the same catalytic domain. To confirm that the signals are from RD21 and e.g. not from a PLCP that is induced upon RD21 expression, we took advantage of a mutant RD21 that carries an additional *N*-glycosylation site in the catalytic domain (**Figure 5A**). This RD21+ mutant runs at a slightly higher molecular weight (MW), caused by the additional *N*-glycan (Gu et al., 2012). Labeling on transiently expressed RD21+ with the probe cocktail revealed fluorescent signals that are similar in intensity and color when compared to wild-type RD21 but they run at a higher MW, consistent with their *N*-glycosylation (**Figure 5**).

**Figure 5.**
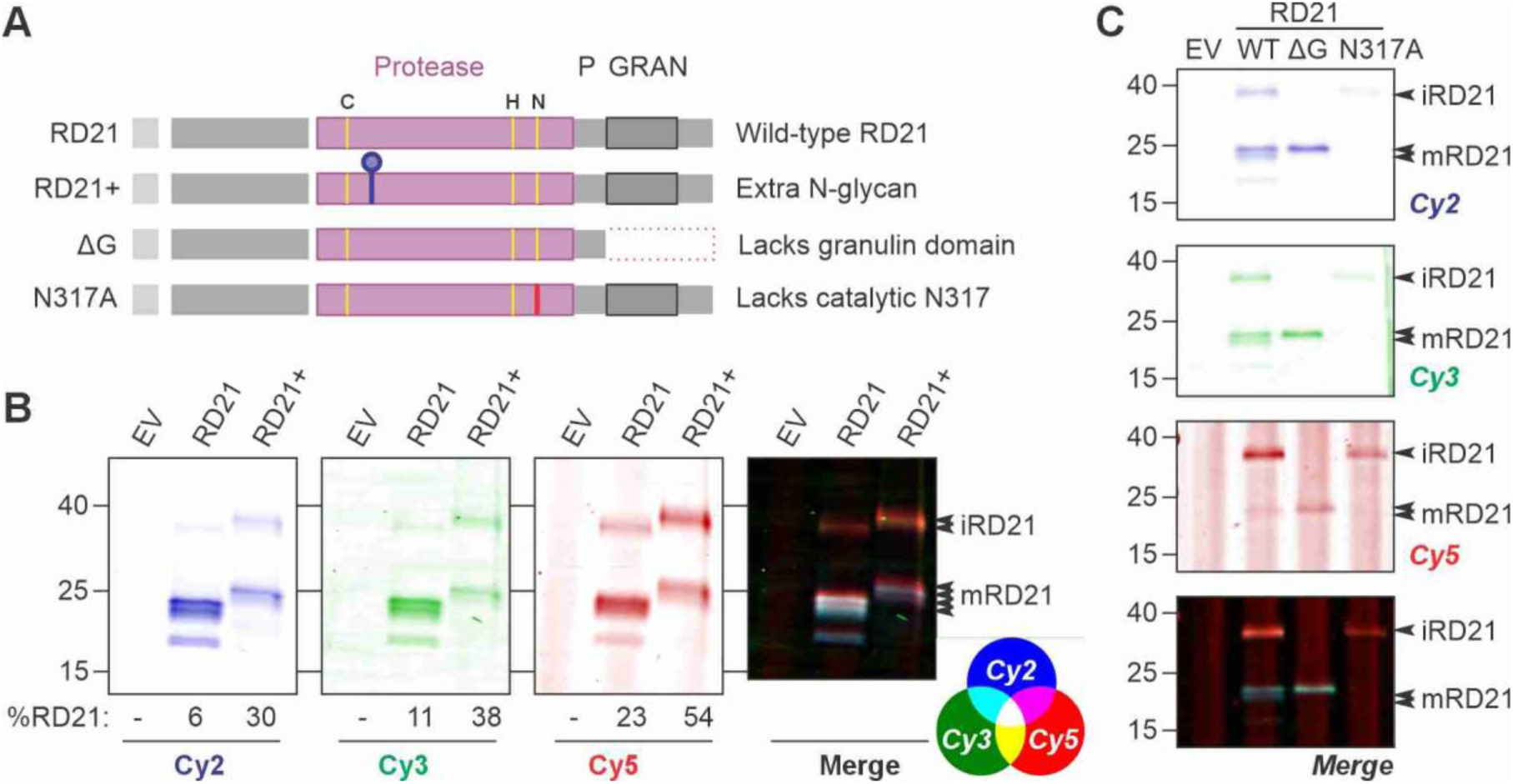
RD21 mutants confirm differential isoform labeling. **(A)** Summary of the used mutants, expressed by agroinfiltration. **(B)** Glycan shift confirms differential isoform labeling of RD21. Extracts of leaves transiently expressing RD21 and the D180N+I182T mutant, which carries an additional sequon for *N*-glycosylation (RD21+), were labeled with a cocktail containing 1 µM TK009, TK010 and TK011 for 3 hours. Proteins were separated on protein gel and scanned for fluorescence. This experiment was repeated twice with a similar outcome. The Coomassie-stained gel is shown in Supplemental **Figure S3**. **(C)** Isoform-fixing mutants confirm differential isoform labeling of RD21. Extracts of leaves transiently expressing RD21 and the granulin deletion mutant (ΔG) and a mutant lacking the catalytic N317 (N317A), were labeled with a cocktail containing 1 µM TK009, TK010 and TK011 for 3 hours. Proteins were separated on protein gel and scanned for fluorescence. This experiment was repeated three times with a similar outcome. The Coomassie-stained gel is shown in Supplemental **Figure S3**.

To further confirm differential labeling of the RD21 isoforms, we took advantage of the ΔG mutant, which lacks the granulin domain, and the N317A mutant, which lacks the catalytic Asp residue N317 and has reduced proteolytic activity and cannot convert itself into mRD21 (Gu et al., 2012, **Figure 5A**). Thus, expression of the ΔG mutant will only produce mature isoform mRD21, whilst the N317A mutant will only produce the intermediate isoform iRD21. Labeling extracts of leaves transiently expressing these two mutants with the probe cocktail resulted in fluorescent signals that was very similar to that seen with the isoforms produced by wild-type RD21: N317A is more efficiently labeled with TK009, similar to iRD21, whilst ΔG is more efficiently labeled by TK010 and TK011, similar to mRD21 (**Figure 5C**). In conclusion, the probes label RD21 isoforms with different efficiencies.

To determine the competitive strength of each probe, we compared the label-and-mix (L&M) and mix-and-label (M&L) intensities of the probes pairwise. Again, the presence of TK011 reduces labeling of both iRD21 and mRD21 isoforms with TK009 and TK010 (**Figure 6A**), demonstrating that TK011 is labeling RD21 faster than both TK009 and TK010. We did not detect this suppression by TK009 on TK010 or TK010 on TK009 labeling (**Figure 6A**), indicating that these two probes have similar competiveness for labeling RD21. Thus, during labeling with probe cocktails, the competitive strength for RD21 labeling is: TK011 > TK010 ∼ TK009.

**Figure 6.**
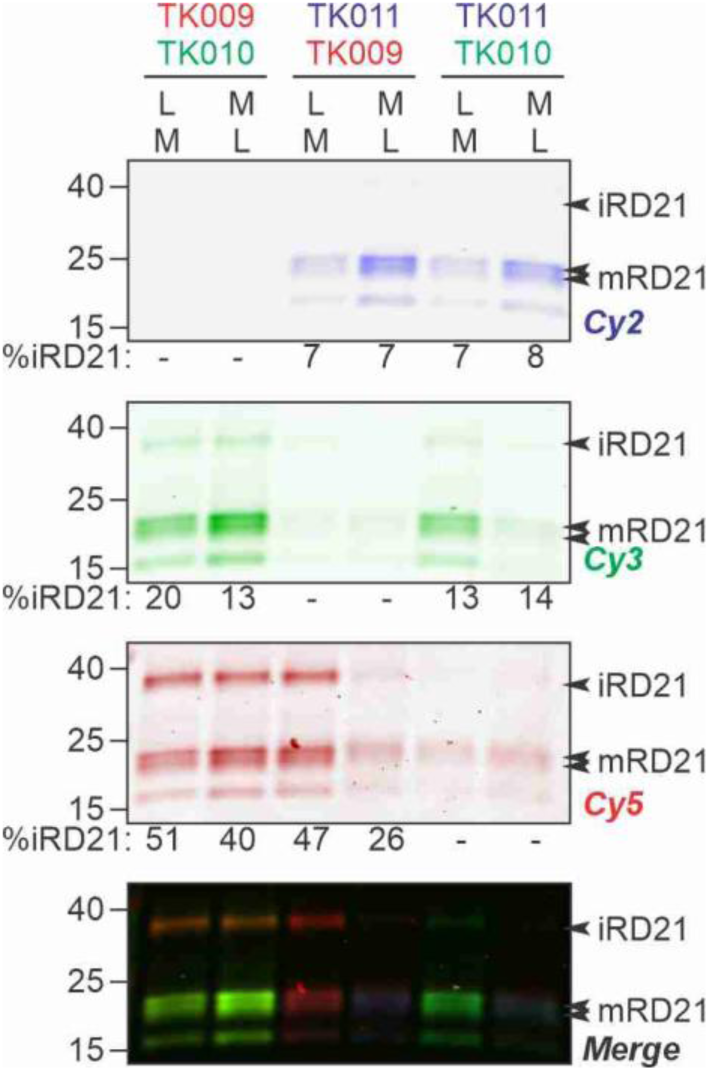
Pairwise competitive labeling reveals relative competitive powers. Extracts of leaves transiently expressing RD21 were labeled with 1 µM TK009, TK010 and TK011 for 3 hours and then mixed (L&M), or labeled with a cocktail containing 1 µM TK009, TK010 and TK011 for 3 hours (M&L). Proteins were separated on protein gel and scanned for fluorescence. This experiment was repeated once with a similar outcome. The Coomassie-stained gel is shown in Supplemental **Figure S3**.

The fact that mRD21 and iRD21 have different probe preferences indicates that the granulin domain in iRD21 influences probe binding in the active site. To investigate this, we generated structural models of iRD21 with Chai-1 (Boitreaud et al., 2024) and found that the orientation of the granulin domain relative to the protease domain can vary because of a flexible Pro-rich linker between the two domains (**Figure 7A**). We next included the probes to generate models of iRD21-probe complexes and found that the granulin domain can act as a lid on the substrate binding groove and directly interact with the fluorophore of the probe (**Figure 7B**). The ability of the granulin domain to flank the substrate binding groove might explain why iRD21 is labeled differently by the probe cocktail than the mRD21 isoform. Because it is longer, the fluorophore of TK009 can reach deeper into the interface pocket, explaining the preferential labeling of iRD21 with this probe.

**Figure 7.**
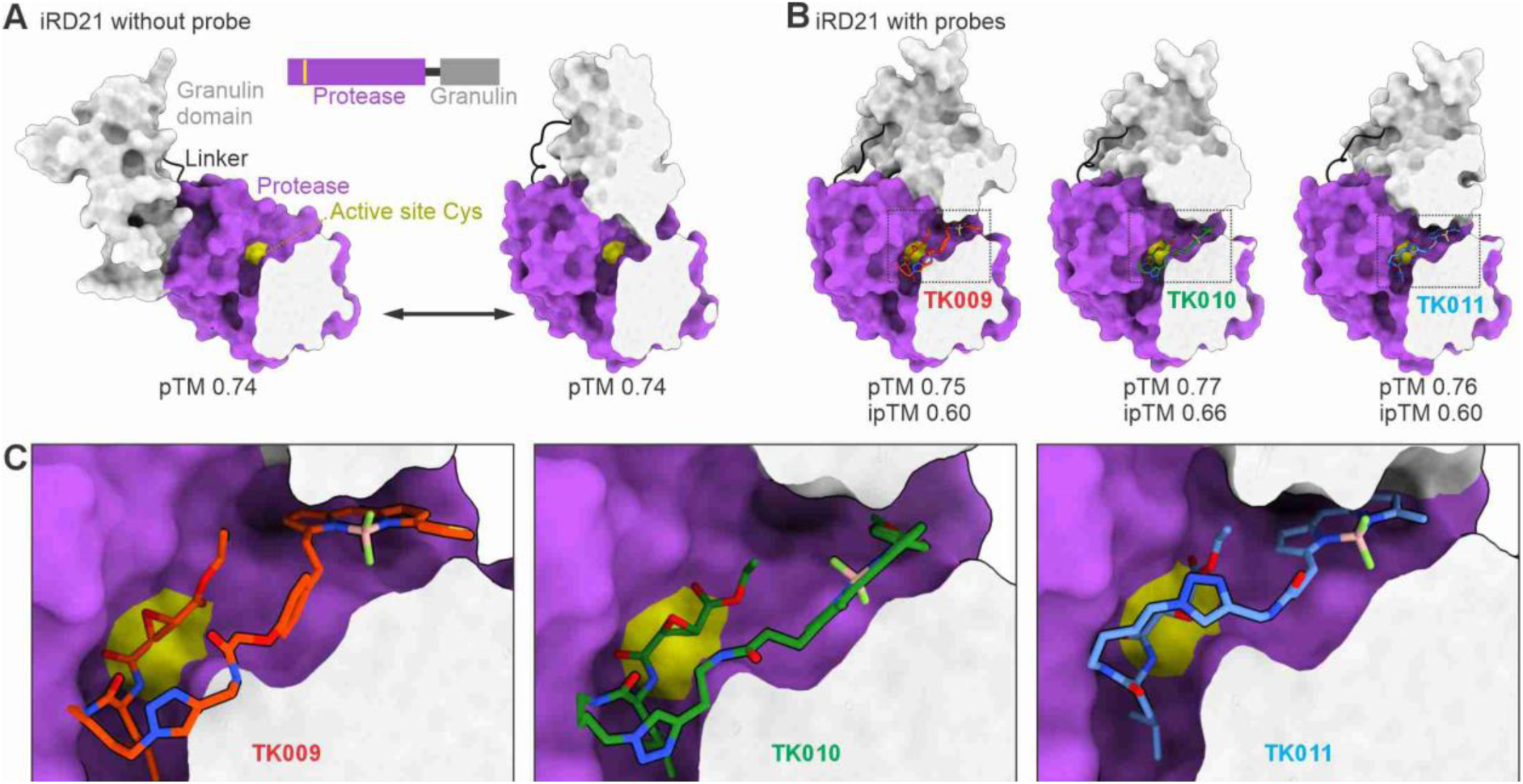
Granulin domain of iRD21 can influence probe binding. **(A)** Structural models of iRD21 without probe, showing two top scoring models adopting different conformations of the granulin domain relative to the protease domain, suggesting a possible flexibility as the two domains are connected via a disordered linker peptide. **(B)** Structural models of iRD21 with probes (TK009, TK010, TK011), showing probe binding in the active site with the granulin domain also forming part of the pocket that can interact with the fluorophore portion of the probe. **(C)** Zoomed in view of the enzyme active site pocket regions (dotted rectangle) of each structure in (B). (A-C) Structural models of iRD21 with or without probes were generated with Chai-1 (Boitreaud et al., 2024) and visualized with ChimeraX (Pettersen et al., 2021). pTM and ipTM scores of each prediction are shown underneath each model. iRD21 structure consists of the granulin domain (grey) connected via the disordered linker peptide (black) to the protease domain (purple) that contain the catalytic cysteine (Cys, yellow) in the active site pocket. Protein structures are displayed as surface. For clarity of the active site pocket view, protein structures are cross-sectioned, with the surface of a clipping plane capped in light grey. Probes are displayed as sticks of different colors (TK009 [red], TK010 [green], TK011 [blue]) with colored heteroatoms (oxygen [dark red], nitrogen [dark blue], fluorine [light green], boron [light pink]).

To test if different PLCPs have different labeling efficiencies for the probe cocktail, we labeled extracts of leaves transiently expressing various PLCPs with the probe cocktail. In addition to RD21, also the mature isoforms of its closely-related paralog RD21B, and its tomato ortholog C14 and *N. benthamiana* ortholog *Nb*RD21 all appeared blue, whilst their intermediate isoforms all appeared red (**Figure 8**), indicating that the granulin domain contributes to substrate selection for all RD21 homologs. The apparent color of Arabidopsis RD21C, however, is distinct from other RD21 family members. Interestingly, PLCPs from other subfamilies appeared with different colors, indicating that subfamilies are labeled with specific polychrome colours. Arabidopsis aleurain-like protease ALP, for instance, appeared yellow because it was efficiently labeled by both TK009 and TK010, whereas RD19A appeared red, caused by strong TK009 labeling, and RD21C, THI1, RD19B and CTB3 appeared pink, caused by labeling with all three probes (**Figure 8**). These experiments indicate that labeling with the probe cocktail results in distinct apparent colors caused by slightly different affinities for the three different probes.

**Figure 8.**
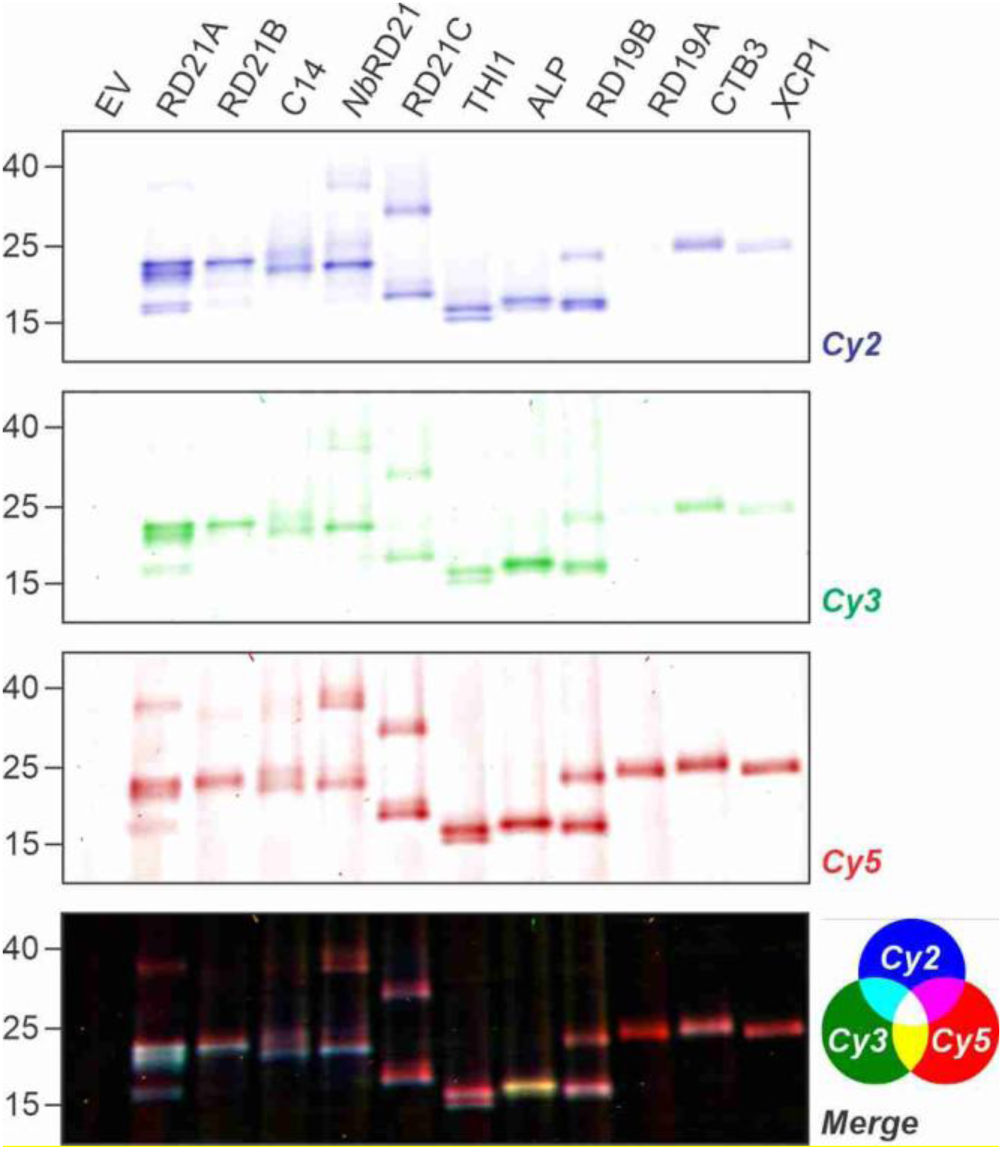
Polychrome labeling distinguishes different proteases. Extracts of leaves transiently expressing various proteases were labeled with a cocktail containing 1 µM TK009, TK010 and TK011 for 3 hours. Proteins were separated on protein gel and scanned for fluorescence. This experiment was repeated twice with a similar outcome. The Coomassie-stained gel is shown in Supplemental **Figure S3**.

Because ALP is distinctly labeled with TK009 and TK010, causing a yellow apparent color (**Figure 8**), we also investigated ALP-probe complexes predicted by Chai-1. ALP is an alaurain-like protease, which are typified by a minichain that originates from the prodomain and is retained in the substrate binding groove with a disulfide bridge that is unique to aleurain proteases (Dodt & Reichwein, 2003). The obstruction of one half of the substrate binding groove is why these proteases process proteins only at the N-terminus as aminopeptidases. Modeling of the minichain-ALP complex places the minichain at the right position (**Figure 9A**). Interestingly, models of minichain-ALP-probe complexes suggest that the minichain may have different conformations and that the fluorophore can interact with the hydrophobic bottom of the peptide binding groove (yellow in **Figure 9B**). This is the same surface that cystatins target with a conserved Trp(W) (Björk et al., 1996). The fluorophores of TK009 and TK010 have larger ring structures with electrons that can create pi-bonds at this interface, when compared to TK011, explaining why ALP has low affinity to TK011.

**Figure 9.**
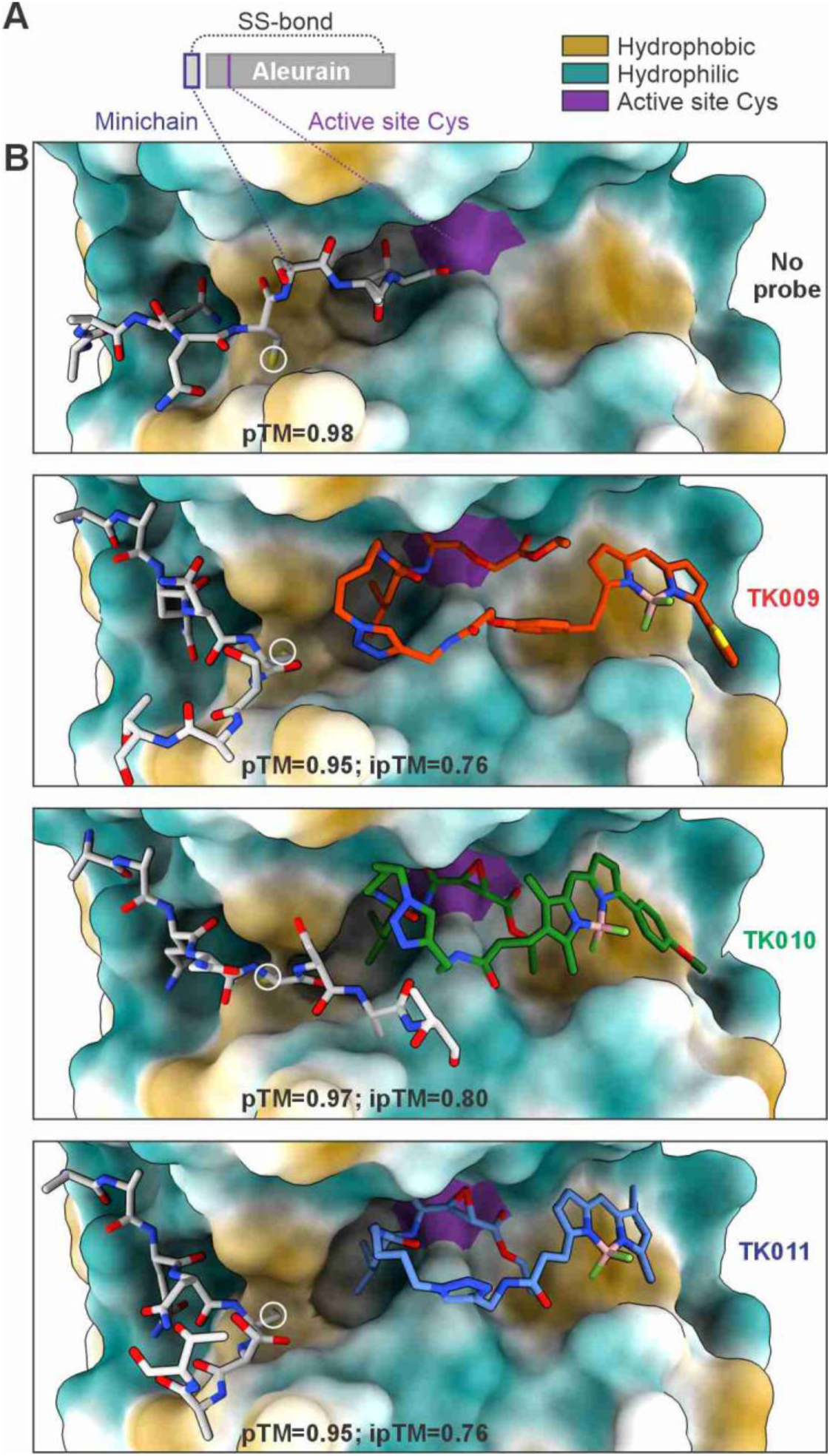
Molecular model of labeling of aleurain-like protease ALP. **(A)** Diagram showing components of mature ALP. **(B)** Structural models of ALP with or without probes, showing probe binding in the active site groove with a hydrophobic patch that can interact with the fluorophore portion of the probes. Structural models were generated with Chai-1 (Boitreaud et al., 2024) and visualized with ChimeraX (Pettersen et al., 2021). pTM and ipTM scores of each prediction are shown underneath each model. The mature ALP structure consists of the aleurain domain (displayed as surface) and the minichain (displayed as stick in light grey), a short peptide occupying part of the active site groove and connected to the protease via a disulfide bond (white circle). Protein surfaces are colored according to hydrophobicity potential (hydrophobic [yellow], hydrophilic [cyan]) and the catalytic cysteine residue (purple). Probes are displayed as sticks of different colors (TK009 [red], TK010 [green], TK011 [blue]) with colored heteroatoms (oxygen [dark red], nitrogen [dark blue], fluorine [light green], boron [light pink]).

Our serendipitous discovery that different probe targets can be distinguished upon competitive labeling with probe cocktails can have important practical implications. We found that different proteases can be distinguished because they react with the probe cocktail with different probe ratios, due to their different affinities to these probes, causing different spectra for each protease and each isoform. Since the probes only differ in their fluorophores, we conclude that these must interact with the protease. Indeed, molecular modeling confirmed that this is possible. These interactions may be less relevant with longer linkers. Interestingly, however, labelling of rat liver lysates with activity-based probes carrying different fluorophores on E-64 also showed slightly different labeling profiles (Greenbaum et al., 2002), indicating that this phenomenon is more wide-spread but underexplored. These probe cocktails now enable us to distinguish proteases and their different isoforms based on the apparent color caused by different reactivities to the probe cocktail. This revealed that the granulin domain in RD21 proteases can interact with the substrate binding groove and this will probably have an influence on selective processing of endogenous RD21 substrates.

The concept of polychrome labeling can be easily expanded to other classes of enzymes for which activity-based probes have been generated. One can use short linkers to create interactions with fluorophores. Alternatively one can also mix probes with slightly different binding groups to generate probe cocktails that can be used for polychrome labeling based on differential probe binding affinities. For instance, a mix of residues in fluorescent activity-based probes targeting caspases, paracaspases or metacaspases (Edgington et al., 2012; Strancar et al., 2022; Hachmann et al., 2015) may be able to display the different proteases in different colors. Likewise, a mix of fluorophosponate warheads (Faucher et al., 2020) might be able to establish polychrome labeling of serine hydrolases. Our experiments show how labeling with polychrome probes can be characterized and validated, and demonstrate that these experiments can deliver novel functional information on enzymes.

## METHODS

### Transient expression

*Agrobacterium tumefaciens* strain GV3101 harboring the binary expression construct was grown overnight in LB medium supplemented with 50 μg/mL kanamycin, 50 μg/mL gentamicin, and 25 μg/mL rifampicin at 28°C with shaking at 200 rpm. Bacterial cells were harvested by centrifugation at 3,500 × g for 10 min at room temperature and resuspending in infiltration buffer (10 mM MES, 10 mM MgCl₂, pH5.7) and 100 μM acetosyringone. The final OD₆₀₀ was adjusted to 0.5, and the suspension was incubated at room temperature in the dark for 2–3 hours prior to infiltration.

Four-week-old *Nicotiana benthamiana* plants were used for infiltration. Bacterial suspensions were introduced into the abaxial side of fully expanded leaves using a needleless 1-mL syringe. Infiltrated plants were maintained under standard growth conditions (22°C, 16 h light/8 h dark photoperiod). Leaves were harvested three days post-infiltration for protein extraction.

### Probe synthesis

Probes were synthesized by click chemistry and purified as described previously (Schuster et al., 2024; Sueldo et al., 2024).

### Protein extraction

Leaf tissue (1 g) was collected from agroinfiltrated leaves at 3DPI and ground to a fine powder in liquid nitrogen using a mortar and pestle. The powder was transferred to a 50 mL centrifuge tube and resuspended in 4 mL ultrapure water by vigorous vortexing. The homogenate was incubated on ice for 10 min and subsequently centrifuged at 5,000 rpm for 5 min at 4°C to remove coarse debris. The resulting supernatant was transferred to a 1.5 mL microcentrifuge tube and further clarified by centrifugation at 15,000 rpm for 10 min at 4°C. The final supernatant was collected and used for protein concentration determination and labelling.

### Labeling, gel electrophoresis and scanning

Protein concentrations were determined using the DC™ Protein Assay Kit I (#5000111, Bio-Rad, Hercules, CA, USA) according to the manufacturer’s instructions. Appropriate amounts of protein extract were used for probe labeling. Unless otherwise indicated, labeling reactions were performed in a final volume of 500 μL containing protein extract, 10 mM DTT, 0.2 μM probe, and 50 mM sodium acetate buffer (pH 6.0). Samples were incubated for 3 h at room temperature in the dark. After labeling, proteins were precipitated by the addition of 1 mL ice-cold acetone pre-chilled at −20°C. Samples were vortexed and centrifuged at 15,000 rpm for 10 min at 4°C. The supernatant was discarded, and protein pellets were air-dried for 2 min in a fume hood. Pellets were subsequently resuspended in 50 μL 2× SDS–PAGE loading buffer and heated at 95°C for 8 min.

Unless otherwise indicated, 20 μg total protein was loaded per lane. Protein samples were separated on 12% SDS–PAGE gels and electrophoresed at 180 V in Invitrogen Novex vertical gel tanks. PageRulerTM Prestained Protein Ladder (26616; Thermo ScientificTM, Waltham, MA, USA) was used as the molecular weight marker. Fluorescently labeled proteins were detected using a Typhoon FLA 9000 imaging system (GE Healthcare Life Sciences, Little Chalfont, UK). Default scanner settings for Cy2 (ex488nm/em525BP20nm), Cy3 (ex532nm/em570BP20nm), and Cy5 (ex635nm/em670BP20nm) fluorophores were used to detect signals from probes TK011, TK010, and TK009, respectively. and intensities were quantified using ImageJ. The relative signal intensity of each target band was calculated as the signal intensity of the target band divided by the sum of the signal intensities of all bands and expressed as a percentage.

### Modeling

Protein structures and binding of probes were predicted using Chai-1 (Boitreaud et al., 2024) with default settings, using with multiple sequence alignment (MSA) generated from jackhmmer search, accessed through chaidiscovery.com. Amino acid sequence of each protein and SMILES string of each probe were used as inputs for modeling (**Supplemental Table S1**). The first ranked model of each prediction was used. ChimeraX (Pettersen et al., 2021) was used to visualize the predicted structures.

## ACKNOWLEDGEMENTS

We thank Urzsula Pyzio for excellent plant care and Sarah Rodgers, Caroline O’Brien and Patricia Bowman for technical support.

## Author contributions

RH conceived the project; TK and MK produced the probes; KZ and MS performed the experiments; NS produced and analysed molecular models; RH wrote the manuscript with input from all authors.

---

**Supplemental Table S1.** Used plasmids.

| Plasmid | Description | Reference |
| --- | --- | --- |
| pRH385 | EV (Empty Vector) | van der Hoorn et al., 2005 |
| P19 | P19 (silencing inhibitor) | van der Hoorn et al., 2005 |
| pRH628 | 35S::RD21 (from Arabidopsis) | Gu et al., 2012 |
| pMS48 | 35S::RD21+PGS (from Arabidopsis) | Gu et al., 2012 |
| pMS17 | 35S::RD21(ΔG) | Gu et al., 2012 |
| pMS15 | 35S::RD21(N317A) | Gu et al., 2012 |
| pTP31 | 35S::RD21B (from Arabidopsis) | Richau et al 2012 |
| pTP41 | 35S::C14 (from tomato) | Shabab et al., 2008 |
| pPB34 | 35S::NbRD21 (from *N. benthamiana*) | Godson et al., 2025 |
| pTP32 | 35S::RD21C (from Arabidopsis) | Richau et al., 2012 |
| pTP33 | 35S::THI1 (from Arabidopsis) | Richau et al., 2012 |
| pFK17 | 35S::AALP (from Arabidopsis) | Richau et al., 2012 |
| pTP38 | 35S::RD19B (from Arabidopsis) | Richau et al., 2012 |
| pTP37 | 35S::RD19A (from Arabidopsis) | Richau et al., 2012 |
| pFK16 | 35S::CTB3 (from Arabidopsis) | Richau et al., 2012 |
| pTP45 | 35S::XCP1 (from Arabidopsis) | Richau et al., 2012 |

**Supplemental Table S2.**
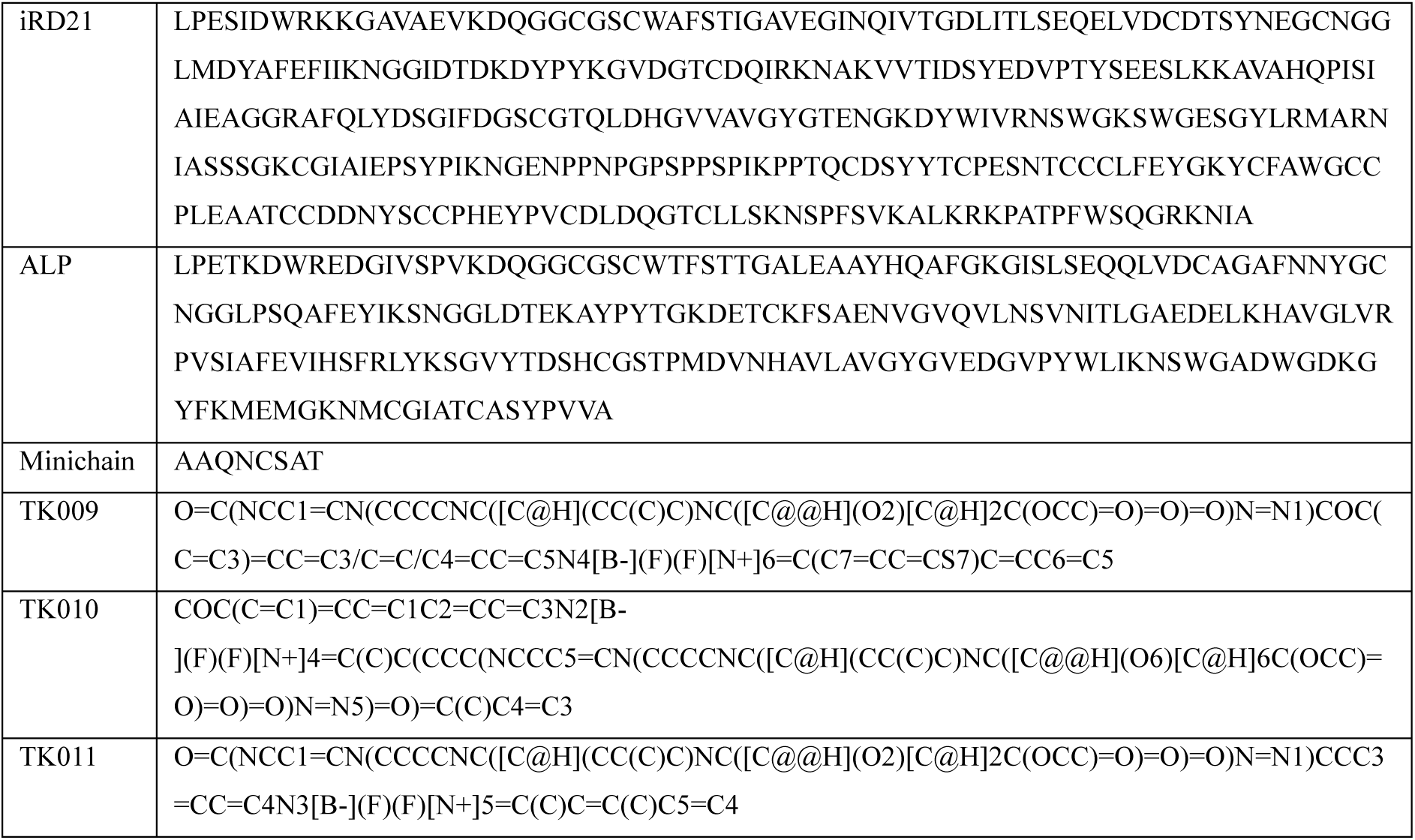
Protein sequences and SMILES strings used for modeling.

## Supplemental Figures

**Figure S1.**
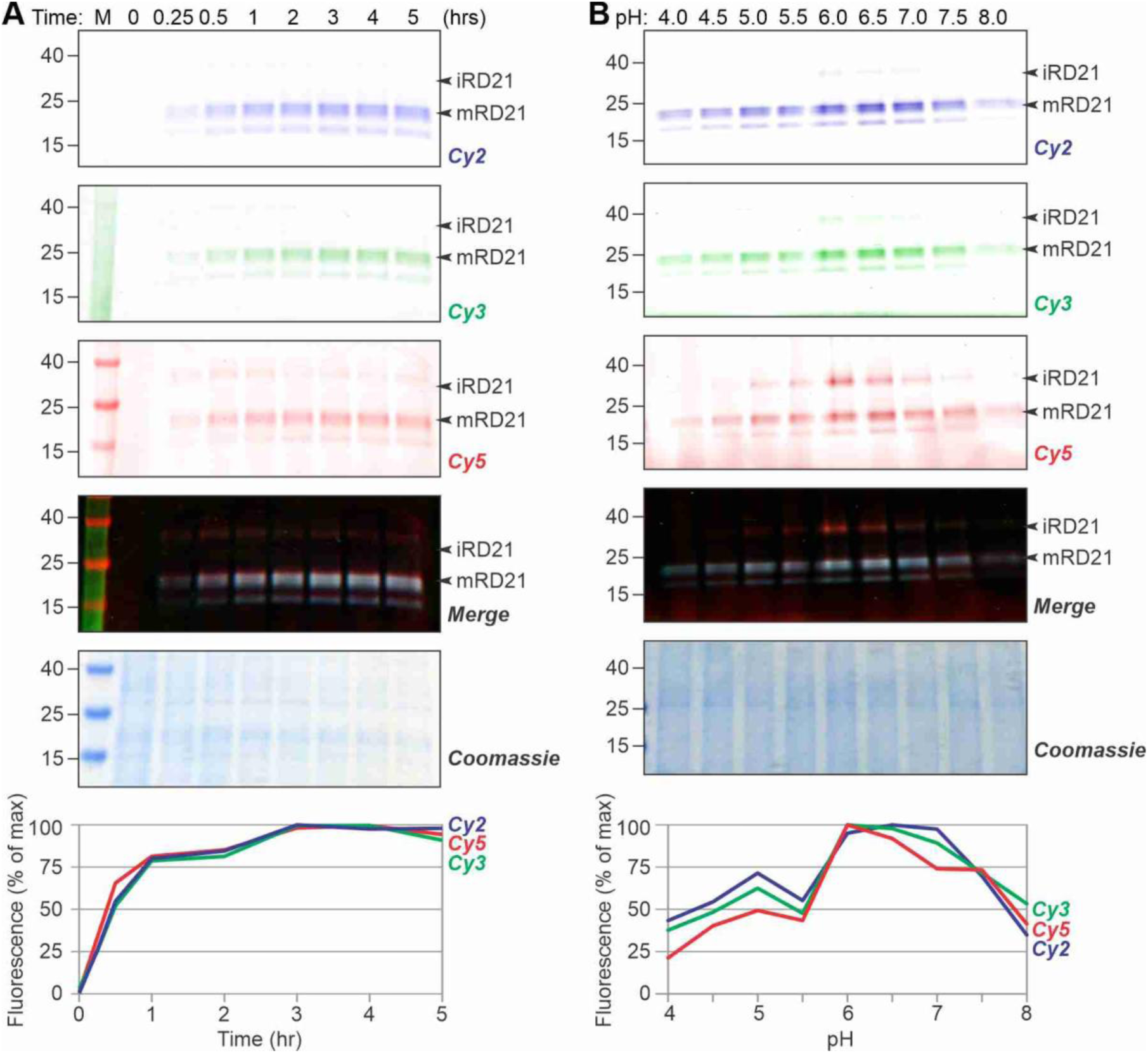
RD21 labeling in time and at various pH. **(A)** Total extracts of agroinfiltrated leaves expressing RD21 were incubated with 1µM TK probe cocktail at pH 6.0 and samples were precipitated in acetone at various timepoints and separated on protein gels, scanned for fluorescence and stained with Coomassie. Fluorescent signal intensities were quantified and plotted as a percentage of the strongest signal for each fluorescent signal. **(B)** Total extracts of agroinfiltrated leaves expressing RD21 were incubated with 1µM TK probe cocktail for 3 hours at various pH and samples were precipitated in acetone and separated on protein gels, scanned for fluorescence and stained with Coomassie. Fluorescent signal intensities were quantified and plotted as a percentage of the strongest signal for each fluorescent signal.

**Figure S2.**
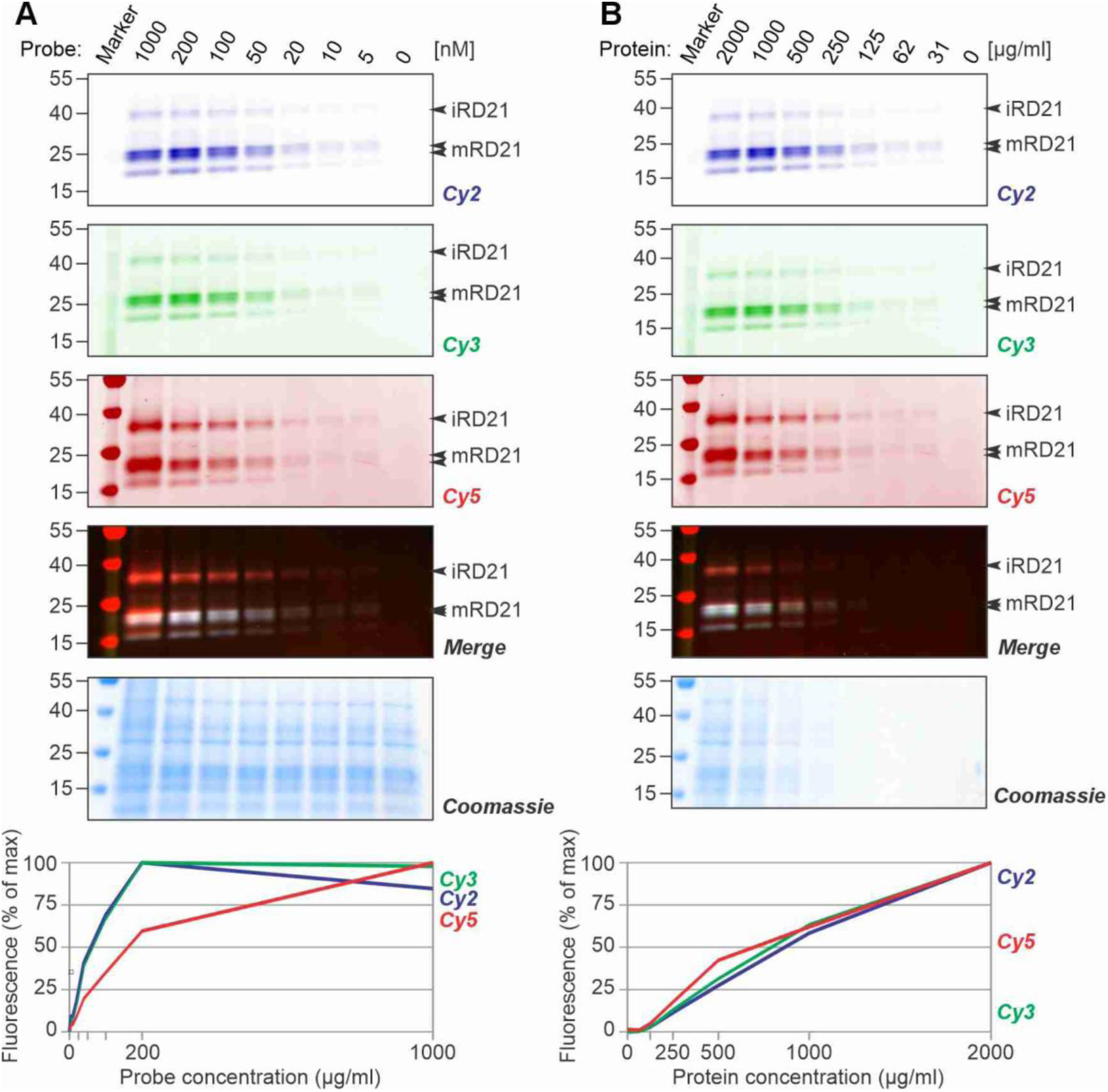
RD21 labeling at various protein and probe concentrations. **(A)** Total extracts of agroinfiltrated leaves expressing RD21 were incubated with various concentrations of TK probe cocktail at pH 6.0 for 3 hours and samples were precipitated in acetone and separated on protein gels, scanned for fluorescence and stained with Coomassie. Fluorescent signal intensities were quantified and plotted as a percentage of the strongest signal for each fluorescent signal. **(B)** Various concentrations of extracts of agroinfiltrated leaves expressing RD21 were incubated with 1µM TK probe cocktail at pH 6.0 for 3 hours and samples were precipitated in acetone and separated on protein gels, scanned for fluorescence and stained with Coomassie. Fluorescent signal intensities were quantified and plotted as a percentage of the strongest signal for each fluorescent signal.

**Figure S3.**
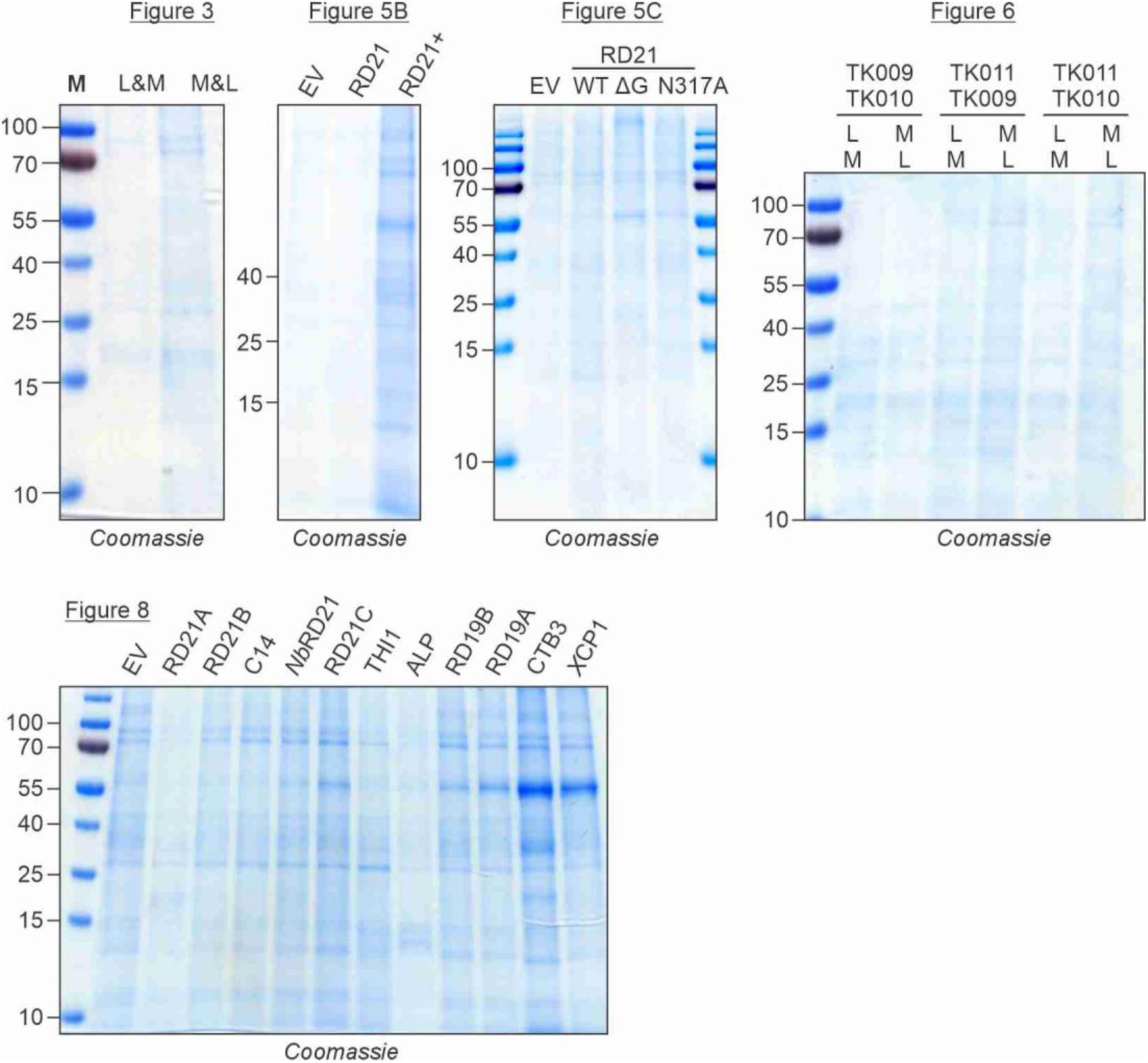
Coomassie-stained gels of labeled samples. Proteomes of protease-expressing leaves were incubated with activity-based probes, separated on protein gels and stained with Coomassie after scanning for fluorescence. Most proteins are degraded by the overexpressed proteases during the 3hr incubation at room temperature without protease inhibitors. More was loaded of the RD21+ sample because the expression was low in this sample.

## REFERENCES

Björk I, Brieditis I, Raub-Segall E, Pol E, Håkansson K, Abrahamson M. (1996) The importance of the second hairpin loop of cystatin C for proteinase binding. Characterization of the interaction of Trp-106 variants of the inhibitor with cysteine proteinases. Biochemistry 35, 10720-10726.

Boitreaud J, Dent J, McPartlon M, Meier J, Reis V, Rogozhnikov A, Wu K (2024) Chai-1: decoding the molecular interactions of life. bioRxiv 2024.10.10.615955.

Chakrabarty S, Kahler JP, van de Plassche MAT, Vanhoutte R, Verhelst SHL. (2019) Recent advances in activity-based protein profiling of proteases. Curr Top Microbiol Immunol. 420, 253–281.

de Bruin G, Xin BT, Kraus M, van der Stelt M, van der Marel GA, Kisselev AF, Driessen C, Florea BI, Overkleeft HS. (2016) A set of activity-based probes to visualize human (immuno)proteasome activities. Angew Chem Int Ed Engl. 55, 4199–203.

Dodt J, Reichwein J. (2003) Human cathepsin H: deletion of the mini-chain switches substrate specificity from aminopeptidase to endopeptidase. Biol Chem. 384, 1327–1332.

Edgington LE, van Raam BJ, Verdoes M, Wierschem C, Salvesen GS, Bogyo M. (2012) An optimized activity-based probe for the study of caspase-6 activation. Chem Biol. 19, 340–352.

Faucher F, Bennett JM, Bogyo M, Lovell S. (2020) Strategies for tuning the selectivity of chemical probes that target serine hydrolases. Cell Chem Biol. 27, 937–952.

Fonović M, Turk B. (2014) Cysteine cathepsins and extracellular matrix degradation. Biochim Biophys Acta. 1840, 2560–2570.

Gillet LC, Namoto K, Ruchti A, Hoving S, Boesch D, Inverardi B, Mueller D, Coulot M, Schindler P, Schweigler P, Bernardi A, Gil-Parrado S. (2008) In-cell selectivity profiling of serine protease inhibitors by activity-based proteomics. Mol Cell Proteomics 7, 1241–1253.

Greenbaum D, Baruch A, Hayrapetian L, Darula Z, Burlingame A, Medzihradszky KF, Bogyo M. (2002) Chemical approaches for functionally probing the proteome. Mol Cell Proteomics. 1, 60–68.

Greenbaum D, Medzihradszky KF, Burlingame A, Bogyo M. (2000) Epoxide electrophiles as activity-dependent cysteine protease profiling and discovery tools. Chem Biol. 7, 569–581.

Gu C, Shabab M, Strasser R, Wolters PJ, Shindo T, Niemer M, Kaschani F, Mach L, van der Hoorn RAL (2012) Post-translational regulation and trafficking of the granulin-containing protease RD21 of *Arabidopsis thaliana*. PLoS One 7, e32422.

Hachmann J, Edgington-Mitchell LE, Poreba M, Sanman LE, Drag M, Bogyo M, Salvesen GS. (2015) Probes to monitor activity of the paracaspase MALT1. Chem Biol 22, 139–147.

Hong TN, van der Hoorn RAL. (2016) DIGE-ABPP by click chemistry: pairwise comparison of serine hydrolase activities from the apoplast of infected plants. Methods Mol Biol. 1127, 183–194.

Hook V, Funkelstein L, Wegrzyn J, Bark S, Kindy M, Hook G. (2012) Cysteine cathepsins in the secretory vesicle produce active peptides: Cathepsin L generates peptide neurotransmitters and cathepsin B produces beta-amyloid of Alzheimer’s disease. Biochim Biophys Acta. 1824, 89–104.

Husaini AM, Morimoto K, Chandrasekar B, Kelly S, Kaschani F, Palmero D, Jiang J, Kaiser M, Ahrazem O, Overkleeft HS, van der Hoorn RAL. (2018) Multiplex fluorescent, activity-based protein profiling identifies active α-glycosidases and other hydrolases in plants. Plant Physiol. 177, 24–37.

Kasperkiewicz P, Altman Y, D’Angelo M, Salvesen GS, Drag M. (2017) Toolbox of fluorescent probes for parallel imaging reveals uneven location of serine proteases in neutrophils. J Am Chem Soc. 139, 10115–10125.

Kocks C, Maehr R, Overkleeft HS, Wang EW, Iyer LK, Lennon-Dumenil AM, Ploegh HL, Kessler BM. (2003) Functional proteomics of the active cysteine protease content in Drosophila S2 cells. Mol Cell Proteomics. 2, 1188–1197.

Kovács J, van der Hoorn RAL. (2016) Twelve ways to confirm targets of activity-based probes in plants. Bioorg Med Chem. 24, 3304–3311.

Misas-Villamil JC, van der Burgh AM, Grosse-Holz F, Bach-Pages M, Kovács J, Kaschani F, Schilasky S, Emon AE, Ruben M, Kaiser M, Overkleeft HS, van der Hoorn RAL. (2017) Subunit-selective proteasome activity profiling uncovers uncoupled proteasome subunit activities during bacterial infections. Plant J. 90, 418–430.

Morimoto K, van der Hoorn RAL. (2016) The increasing impact of activity-based protein profiling in plant science. Plant Cell Physiol. 57, 446–461.

Niphakis MJ, Cravatt BF. (2024) Ligand discovery by activity-based protein profiling. Cell Chem Biol. 31, 1636–1651.

Passarge A, Demir F, Green K, Depotter JRL, Scott B, Huesgen PF, Doehlemann G, Misas Villamil JC. (2021) Host apoplastic cysteine protease activity is suppressed during the mutualistic association of *Lolium perenne* and *Epichloë festucae*. J Exp Bot. 72, 3410–3426.

Pettersen EF, Goddard TD, Huang CC, Meng EC, Couch GS, Croll TI, Morris JH, Ferrin TE. (2021) UCSF ChimeraX: Structure visualization for researchers, educators, and developers. Protein Sci. 30, 70–82.

Rawlings ND, Barrett AJ, Thomas PD, Huang X, Bateman A, Finn RD. (2018) The MEROPS database of proteolytic enzymes, their substrates and inhibitors in 2017 and a comparison with peptidases in the PANTHER database. Nucleic Acids Res 46, D624–D632.

Richau KH, Kaschani F, Verdoes M, Pansuriya TC, Niessen S, Stüber K, Colby T, Overkleeft HS, Bogyo M, Van der Hoorn RAL. (2012) Subclassification and biochemical analysis of plant papain-like cysteine proteases displays subfamily-specific characteristics. Plant Physiol. 158, 1583–1599.

Schulze-Hüynck J, Kaschani F, van der Linde K, Ziemann S, Müller AN, Colby T, Kaiser M, Misas Villamil JC, Doehlemann G. (2019) Proteases underground: analysis of the maize root apoplast identifies organ specific papain-like cysteine protease activity. Front Plant Sci. 10, 473.

Schuster M, Eisele S, Armas-Egas L, Kessenbrock T, Kourelis J, Kaiser M, van der Hoorn RAL. (2024) Enhanced late blight resistance by engineering an EpiC2B-insensitive immune protease. Plant Biotechnol J. 22, 284–286.

Shabab M, Shindo T, Gu C, Kaschani F, Pansuriya T, Chintha R, Harzen A, Colby T, Kamoun S, van der Hoorn RAL. (2008) Fungal effector protein AVR2 targets diversifying defense-related cys proteases of tomato. Plant Cell 20, 1169–1183.

Štrancar V, van Midden KP, Krahn D, Morimoto K, Novinec M, Funk C, Stael S, Schofield CJ, Klemenčič M, van der Hoorn RAL. (2022) Activity-based probes trap early active intermediates during metacaspase activation. iScience 25, 105247.

Sueldo DJ, Godson A, Kaschani F, Krahn D, Kessenbrock T, Buscaill P, Schofield CJ, Kaiser M, van der Hoorn RAL. (2024) Activity-based proteomics uncovers suppressed hydrolases and a neo-functionalised antibacterial enzyme at the plant-pathogen interface. New Phytol. 241, 394–408.

van der Hoorn RAL, Leeuwenburgh MA, Bogyo M, Joosten MHAJ, Peck SC. (2004) Activity profiling of papain-like cysteine proteases in plants. Plant Physiol. 135, 1170–1178.

